# Rising temperatures favour parasite virulence and parallel molecular evolution following a host jump

**DOI:** 10.1101/2025.01.14.632940

**Authors:** Tobias E. Hector, Julia M. Kreiner, James C. Forward, Kim L. Hoang, Emily J. Stevens, Serena Johnson, Jingdi Li, Kayla C. King

**Affiliations:** Department of Biology, University of Oxford; Oxford, UK, OX1 3SZ; Biodiversity Research Centre, University of British Columbia; Vancouver, BC Canada V6T 1Z4; Department of Ecology & Evolution, University of Chicago; IL 60637, United States; Department of Microbiology & Immunology, University of British Columbia; Vancouver, BC Canada; Emory School of Medicine, Emory University; Atlanta, GA 30322, United States; School of Life Sciences, Keele University; Staffordshire, UK, ST5 5BG; Department of Zoology, University of British Columbia; Vancouver, BC Canada

## Abstract

Climate change is facilitating the poleward emergence of parasites, increasing the risk of jumping into new animal species, including humans. Whether more virulent or transmissible variants will spread during these climate-driven outbreaks is unclear. We experimentally evolved a wild parasitic bacterium, across the thermal range (20-30°C) and extremes (35°C) of Cape Verde – the site of field collection – in a novel, temperate animal host. At the parasite’s typical warm environmental temperature, we found that virulence escalated across evolutionary time. Parasites evolved at hot temperatures, towards the limit of host-parasite survival, displayed a ‘cryptic’ virulence boost, deadlier once infecting animals at cooler temperatures. Patterns of molecular evolution were constrained to parallel changes in fewer loci at extreme temperatures. Our findings suggest that rising temperatures will leave predictable phenotypic and genomic signatures on evolving parasites as they emerge with climate change.

## Main Text

Climate change is associated with outbreaks of emerging infectious diseases (*1, 2*). As temperate regions warm and become suitable for parasites from lower latitudes (*3*–*5*), there are heightened risks of jumps into temperate host species (*6, 7*). The resulting mismatches between host and parasite thermal tolerances are predicted to exacerbate infections when warm temperatures are favourable for the parasite (*5*). Yet, when parasites colonise a novel species, they cannot fully exploit host resources. This suboptimal environment forces parasites to adapt quickly, sometimes losing or gaining virulence (i.e., harm caused to infected hosts) to improve ongoing transmission (*8*–*10*). The dynamics and outcomes of parasite evolution are understudied in most wild, emerging disease systems (*11, 12*). Local and global outbreaks of zoonotic diseases have revealed an urgent need to forecast evolutionary trajectories of virulence following a host jump (*13, 14*), particularly in an era now defined by warming and extreme temperatures (*15*).

It remains unclear to what extent temperature can shape parasite evolution (*16, 17*). When not at evolutionary equilibrium (*18, 19*), virulence evolution can become decoupled from replication and transmission (*20, 21*), particularly if initial parasite adaptation occurs in traits not directly involved in host exploitation (*e*.*g*., immune evasion) (*22*). For tropical parasites emerging into temperate host species (*5, 6*), warming might favour high virulence if parasite fitness is not optimal in the new environment (*14*) or if host tolerance is enhanced (*23*). An alternative trajectory might arise when a host’s ability to withstand heat is impaired by infection (*24, 25*). Warmer hosts being sicker might shorten the infectious period – an outcome predicted to either favour less virulent variants (*17, 26*) or constrain adaptive evolution due to population bottlenecks (*27*). Temperature is also a key selective force on parasite free-living stages. Warming can extend the seasonal window for parasite growth, directly contributing to their development and between-host transmission (*28*–*30*), but can reduce the survival of some free-living stages in the environment (*29, 31*). Virulence evolution will ultimately depend on the relative strength and direction of selection, as well as the trade-offs involved in adapting to the within- and between-host environments as temperatures climb (*32, 33*).

To test these predictions, we experimentally evolved a wild bacterial parasite (*Leucobacter musarum*) after introduction to *Caenorhabditis elegans* nematodes at ecologically relevant temperatures. *Leucobacter* species have a global distribution and are frequently found as animal associates (including with *Caenorhabditis* nematodes) in natural and agricultural systems (*34*). The parasite was isolated from *C. tropicalis* (strain JU1635) in rotten banana pseudostems in Cape Verde (Fig. 1A) (*35*). Cape Verde is a tropical archipelago in the central Atlantic Ocean, where annual average surface soil temperatures are around 25°C, ranging between 19°C and 28°C, with maximum average soil temperatures as high as 36°C (Fig. 1B) (*36*). During infection in the widespread *C. elegans* (Fig. 1A) *–* specifically, the canonical N2 isolated from Bristol, England – *L. musarum* attaches to the cuticle before infiltrating the host via the rectum, rapidly inducing systemic infection, reduced fecundity, and death (Fig. 1C and fig. S1) (*35*). In its sympatric *C. tropicalis* host isolate, *L. musarum* is sublethal, but reduces reproductive output (Fig. 1C and fig. S1). As with other emerging parasites, infection outcomes are more severe for the novel host than in host populations with a history of previous exposures and opportunities for an evolutionary response (*10, 11, 37, 38*).

**Fig. 1.**
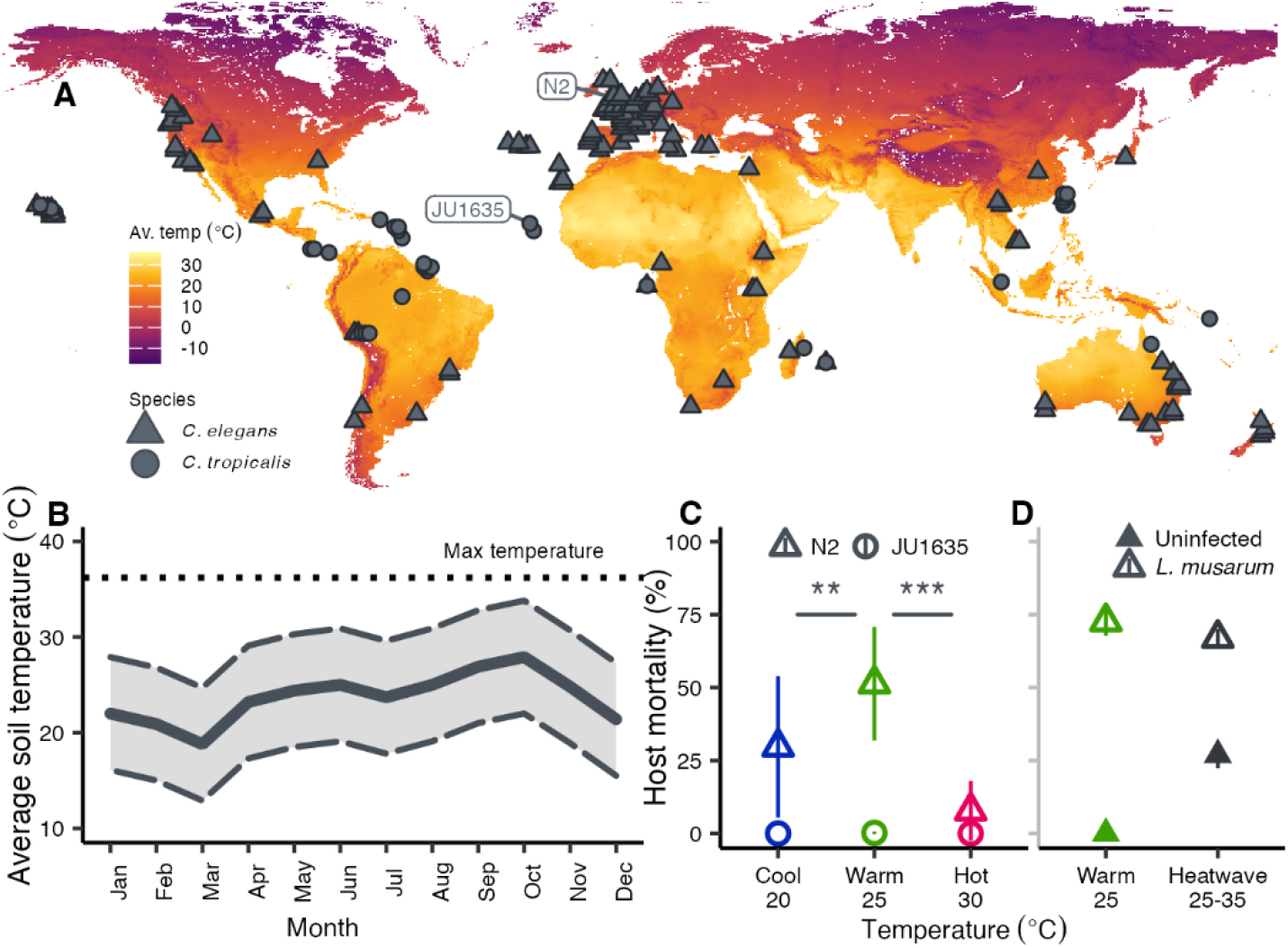
Temperature natural history of *Caenorhabditis* nematodes and the associated virulence of bacterial parasite *Leucobacter musarum*. A) Known distributions of the natural host (*C. tropicalis*) and the widespread novel host (*C. elegans*) for *L. musarum* [data from (*45, 46*)]. Colours represent annual average soil surface temperatures (*36*). Collection locations for *C. tropicalis* JU1635 from which *L. musarum* was isolated and the novel *C. elegans* N2 hosts are labelled. B) Predicted average soil temperatures (at 0.5cm depth) for each month (solid line), average diurnal variation (dashed lines), and the average maximum soil temperature (dotted line) from the location *L. musarum* was isolated [data from (*36*)]. C) *C. elegans* N2 and *C. tropicalis* JU1635 mortality when exposed to the ancestral *L. musarum* clone at three ecologically relevant constant temperatures (Supp. Methods). Mortality is low for *C. tropicalis*, but reproduction is severely limited by infection and hot temperatures (fig. S1). Uninfected *C. elegans* have negligible mortality across this thermal range. Stars show significant pairwise comparisons between both 20°C (Dunnet’s post hoc test, z = −3.485, *P* = 0.001) and 30°C (Dunnet’s post hoc test, z = −11.56, *P* = <0.001) relative to 25°C for infected *C. elegans* N2 hosts. Mean values (± 1 SD) are shown. D) Simulated heatwaves (four-hour ramp between 25°C and 35°C) cause mortality in *C. elegans* populations, particularly when exposed to *L. musarum* (Tukey’s post hoc test comparing infected and uninfected mortality under heatwave, t=15.202, p=<.0001).

Parasite-induced host mortality – a proxy for virulence – peaks during infections at the parasite’s typical warm environmental temperature (25°C; Fig. 1C) (betabinomial ANOVA, χ^2^ = 135.57, d.f. = 2, *P* = < 0.001). *Caenorhabditis elegans* are normally reared at 15-20°C and are stressed at 25°C with reduced fecundity (fig. S1) (*39, 40*). At 20°C (cool), *L. musarum* is less virulent to this temperate host (Fig. 1C). Virulence declines quickly during infections at hot environmental temperatures (30°C, Fig. 1C). Despite a trend for increased *in vitro* growth at 30°C, replication within hosts (parasite load, e.g. (*41*)) remains relatively stable across the constant temperature range (fig. S2) (Kruskal-Wallis Test χ^2^ = 5.956, d.f. = 2, *P* = 0.051; Dunn post hoc test 25°C-30°C, z = 2.385, *P* = 0.017, adj. *P* = 0.051). By activating immune pathways (*42*) and inducing structural changes in the cuticle (*43*), hot conditions can promote tolerance to parasites. So, hotter hosts are not necessarily sicker. Towards the peak temperatures in Cape Verde, and increasingly in temperate regions where *C. elegans* N2 originates (35°C simulated heatwave; Fig. 2A) (*44*), *L. musarum* again becomes relatively virulent (Fig. 1D) (ANOVA, F_3,12_ = 342.2, *P* = <0.001; infected vs uninfected Tukey’s post hoc test t=15.202, p=<0.001) whilst simultaneously experiencing reduced cell viability relative to at 25°C (fig. S3) (Kruskal-Wallis Test χ^2^=14.552, d.f.=1,p=<0.001). These findings reveal that temperature shifts can alter the nature of the virulence-replication relationship, with heatwaves driving virulent infections despite fitness costs for the parasite (*17*).

**Fig. 2.**
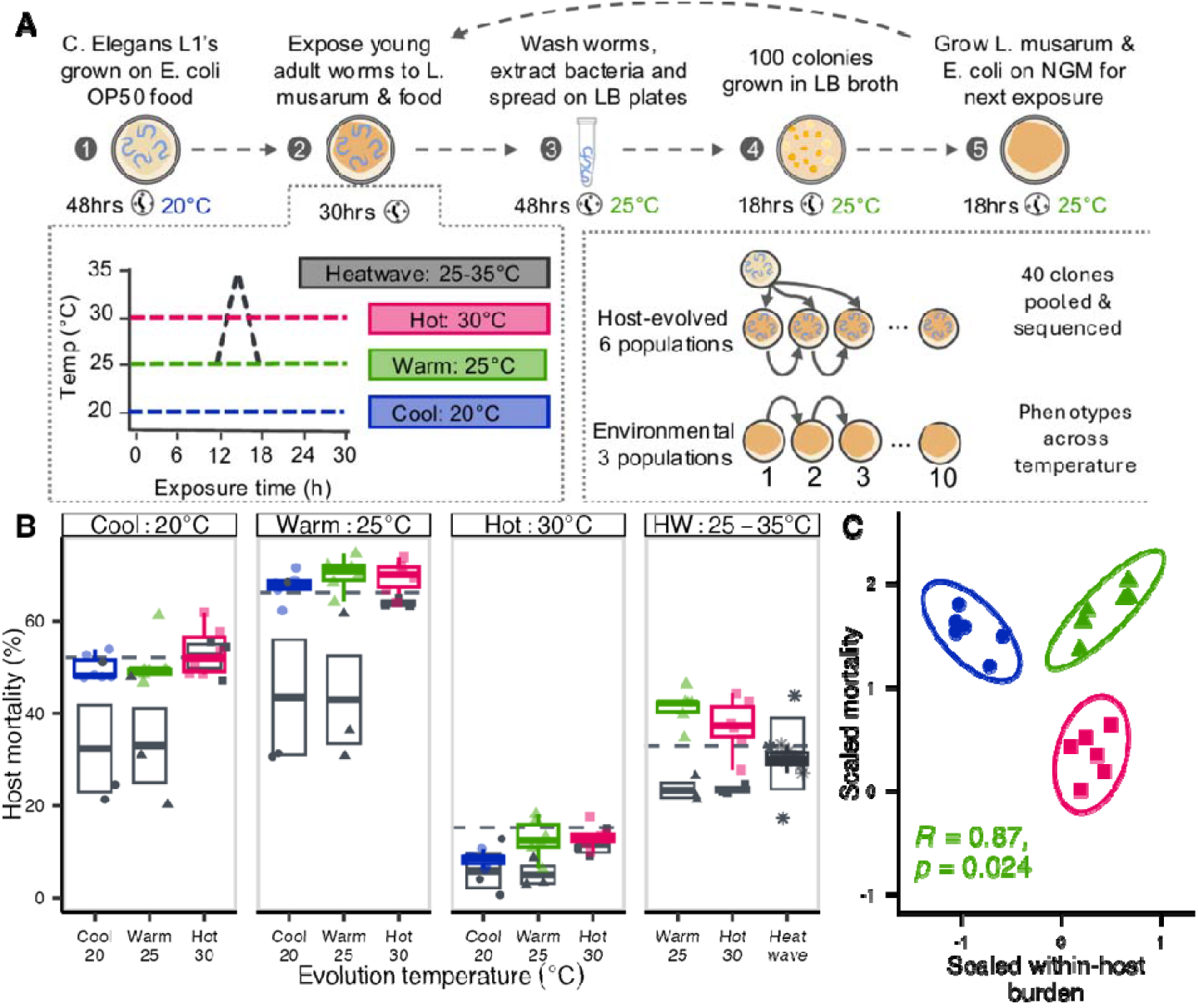
Experimental evolution reveals the impact of temperature on parasite evolution. A) visual overview of experimental evolution protocol. Parasite lineages only experienced differences in temperature during experimental infections, all other *in vitro* culturing took place at either 25°C (for *L. musarum*) or 20°C (for *C. elegans*). We included three ecologically relevant constant temperatures (cool: 20°C, warm: 25°C, and hot: 30°C), and a “heat wave” treatment (HW). During heat wave, temperatures were quickly ramped from 25°C to 35°C (∼0.1°C/min) and then back to 25°C over a 4-hour period. These peak temperatures are beyond the thermal maxima for both the host and parasite (fig. S3). B) Mortality (%) of *C. elegans* hosts exposed to evolved parasite populations (Passage 10) at each temperature (facets show the assay temperature at which mortality was measured) in a factorial common garden experiment. Host-evolved lineages are shown with coloured boxplots and points, and grey boxes and points show the mean and standard deviation of the environmental control populations. Dashed horizontal lines show the mean ancestral phenotype. C) Relationship between within-host bacterial burden and host mortality at 25°C for host evolved lineages (population means centred relative to their respective no-host controls and scaled to allow comparison, colours correspond to those in B), showing Pearson’s correlation across 25°C evolved populations (green).

We experimentally passaged parasite populations independently in nematode populations across four temperature regimes (Fig. 2A). Environmental (host-free) controls were conducted alongside at each temperature. We carried out phenotypic assays of host mortality upon infection (metric for parasite virulence) and load (metric for parasite replication) across treatments. We then used shotgun sequencing of pools of colonies to measure evolutionary changes in the genomic composition of *L. musarum* populations and to identify molecular targets of selection (supp. Methods).

We found that the parasite’s typical warm temperature favoured higher virulence in the novel, temperate host species. Warming at 25°C selected for the highest levels of host mortality during infection, overall and compared to the ancestor (Fig. 2B) (LMM with ancestral virulence as intercept, within-host 25°C t=2.646, df=45, p=0.011, tables S1-4). These parasite populations were also 20-30% deadlier than environmental controls under the same warm temperature (Host/Environmental Log odds ratio=3.11, z= 5.017, p<0.0001), and during subsequent infections across the full thermal range (Fig. 2B; table S4). When passaged at cooler temperatures favourable for the host, ancestral levels of virulence were maintained in host-evolved parasite populations (Fig. 2B) (LMM with ancestral virulence as intercept, within-host 20°C, t=-1.544, df=45, p=0.13; table S2), while environmental controls lost virulence (Host/Environmental Log odds ratio=2.17, z=3.362, p=0.0008). These patterns of virulence were generally consistent across cool and warm assay temperatures (Fig. 2B; table S3) (ANOVA 3-way interaction term, χ^2^=1.152, df= 4, p=0.886), with virulence depending on the interaction between host environment and passage temperature (ANOVA, χ^2^=17.9, df=2, p=<0.001). Our findings suggest that the capacity for virulence to increase following a host jump may be maximised in thermal environments most optimal for the parasite. The threat to host populations from emerging parasites alongside unseasonably warming temperatures (*5, 6*) will likely be amplified by rapid parasite evolution.

The virulence-transmission trade-off hypothesis predicts virulence and replication to be positively correlated in environments optimal for host exploitation (*8, 18, 19*). These parasite traits can thus be decoupled in emerging parasites when host exploitation is suboptimal (*12, 14, 20, 47*). We found that virulence and replication evolved along different trajectories across temperatures (Fig. 2C, fig. S4 and table S5) (MANOVA, interaction between evolution and assay temperature for host-evolved lines: Pillai statistic=0.549, F_8,90_=4.256, p=<0.001). Within-host evolution during warming led to a positive correlation between virulence and replication (Fig. 2C) (Pearson’s *R* = 0.87, p= 0.024), however passage at cooler temperatures towards the host’s optimum, broke down this relationship (Fig. 2C) (20°C, Pearson’s R=-0.57, p=0.24). Our results suggest that temperature can cause these parasite traits to evolve along independent trajectories during thermal mismatches. The predictability of parasite virulence evolution will be context dependent, potentially deviating from expected trajectories at temperatures suboptimal for invading tropical parasites.

Hot temperatures – beyond the host’s thermal maxima – drove ‘cryptic’ evolutionary changes in virulence which were only apparent in different thermal contexts. At sympatric temperatures, parasite virulence was similar to the ancestor and environmental controls following passage at 30°C (host/ancestor t=-1.298, p=0.201; host/Environmental Log odds ratio=1.17, z=0. 610, p=0.542; tables S1-4) and simulated heatwaves (host/ancestor t=-1.249, p=0.225; Host/Environmental Log odds ratio=0.998, z=-0.013, p=0.989; table S6-7) (Fig. 2B). Virulence and replication were not correlated for these populations (Fig. 2C; R = 0.27, p-value = 0.6). Yet, parasites passaged at hot temperatures were more virulent compared to the ancestor or controls when returned to warm conditions (t=2.173, p=0.035; table S2) and under a simulated heatwave (Host/Environmental Log odds ratio=0.513, z=-3,632, p=0.0003; table S6-7) (Fig. 2B). Higher virulence was favoured at constant hot temperatures, an evolutionary outcome predicted when hosts are tolerant to infection (*48, 49*). On average, nematodes could better withstand all evolved parasite lineages during constant hot conditions (Fig. 2B) (host mortality 30°C vs. 20°C: z=-20.736, p=<.0001; 30°C vs. 25°C: z=-26.655, p=<.0001) while maintaining relatively high parasite loads (fig. S5). In *C. elegans*, as in most eukaryotes, heat triggers multiple stress response systems, such as the heat-shock and unfolded protein responses, which can confer either resistance or tolerance to infection (*42, 50, 51*). Ultimately, rising average temperatures towards the upper thermal limits of temperate hosts may encourage variants which are deadlier when host tolerance subsides at lower temperatures or during future heat waves. This evolutionary process may play a role in mass mortality events and seasonal disease outbreaks from bacterial infections observed in wild and farmed animals (*52, 53*), if at the end of unusually hot periods.

Many bacterial parasites can proliferate in the environment. Understanding the impact of temperature on evolutionary trajectories during environmental residency is crucial to forecasts of infection outcomes upon host contact. We introduced parasite populations evolved *in vitro* across temperatures (i.e., environmental controls) to host populations. We found that virulence was rapidly lost following environmental passage at cool and warm temperatures (Fig. 2B; table S4). A positive relationship between *in vitro* lag time (supp. Methods) and virulence (fig. S6) (Pearson’s correlation *R*=0.55, p=0.019) points towards a functional trade-off between environmental growth and virulence. Loss of virulence for environment-adapted parasites also coincided with the lowest rates of biofilm formation (Fig. 3A and fig. S7) (ANOVA, F(5,21)=14.851, p=<0.001; 20C host/environment post hoc: t=-6.518, p=<0.001; 25C host/environment: t=-4.906, p=<0.001; Supp. methods). These results support the prediction that environmental residency can select against costly virulence-related traits (*33*). At hot and heatwave temperatures, however, the virulence of environmental parasites was maintained (Fig. 2B) (host/environment log odds ratio=1.17, z=0.610, p=0.542, table S4 and S7). Consequently, when returned to warm temperatures, hot-evolved parasites remained highly virulent irrespective of their *in vitro* or *in vivo* evolutionary history (log odds ratio=1.32, z=1.220, p=0.223). Highly deadly variants residing in the environment may thus be conserved as temperatures continue to escalate.

**Fig. 3.**
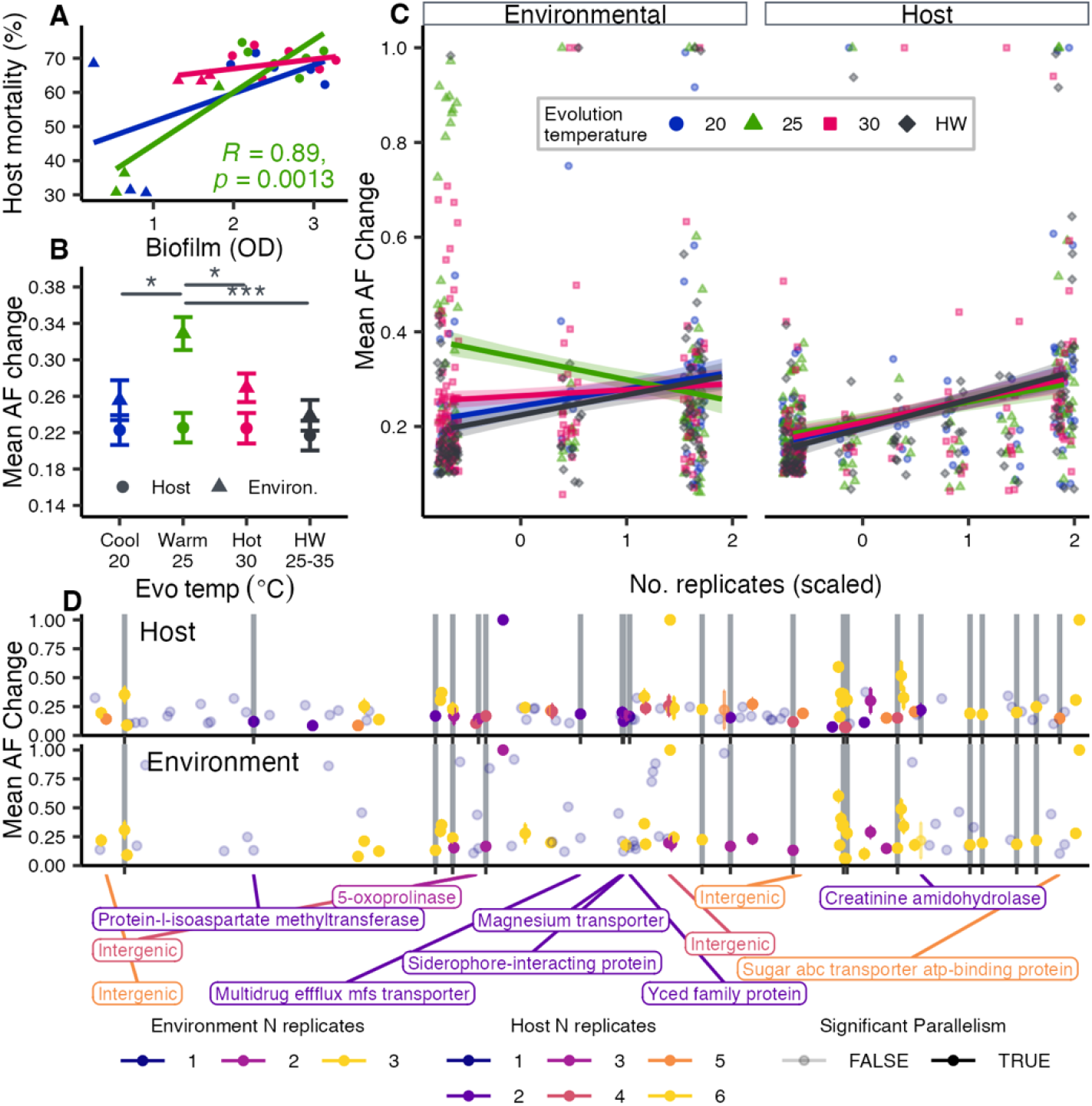
Temperature and hosts drive parasite genomic evolution. A) Spearman’s rank correlation between population mean biofilm formation (optical density (600), see Supp. Methods) and mean host mortality in *C. elegans* exposed to each evolved parasite population in an assay at 25°C. Shown is the significant Pearson’s correlation coefficient for host-evolved and environmental control populations passaged at 25°C. B) Predicted mean (±SE) frequency of *de novo* variants (MAF >0.1 or showing significant parallel change across replicate populations) for different combinations of host and temperature treatment. Stars correspond to post hoc pairwise comparisons between environmental parasites evolved at 25°C and each other temperature (pairwise t-test’s 25°C-20°C: t=-2.689, p=0.021, 25°C-30°C: t=-2.637, p=0.024, and 25°C-HW: t=-3.776, p=<0.001). C) Predicted mean (±SE) frequency of *de novo* variants as a function of the number of replicate populations a SNP arose in parallel (scaled within each host treatment). D) Allele frequency change for *de novo* variants (MAF >0.1 or showing FDR corrected significant parallel change across replicate populations) by genomic position for host-evolved and environmental controls passaged at 25°C. Colours show the level of parallelism across replicate populations (note that host treatments had six replicate populations and environmental controls had three), transparent points did not display significant parallel evolution. Grey bars show parallel variants within genes (synonymous and non-synonymous). Labels below highlight the predicted functions of genes containing variants that were unique to host-evolved replicate populations.

Using pooled whole genome sequencing, we quantified the evolution and putative signals of selection on *de novo* variants arising over ten host generations (Fig. 2A; Supp. methods). Across treatments, there were approximately 70 *de novo* variants per evolved lineage with 4-8 SNPs reaching allele frequencies >50% (fig. S8A; table S8). For parasite populations evolved within hosts, ∼22% of variants were unique to each temperature treatment and 10% were shared across all treatments (fig. S9). Within coding regions, 70% of variants were non-synonymous substitutions (fig. S8B) indicating potential targets of selection, many within genes linked to biofilm formation, virulence, metabolism, and abiotic stress responses (figs. S10-S11). Across warm-adapted parasite populations evolved in hosts and the environment virulence was positively correlated with *in vitro* biofilm formation (Fig. 3A) (Pearson’s correlation R=0.89, p=<0.001), suggesting a polygenic basis to shifts in virulence (*54*).

Parasite molecular evolution across the genome was on average lower, but more predictable, within novel hosts compared to within the environment. When looking at common mutations (MAF > 0.1), within-host evolution generated the lowest genome-wide diversity across the full thermal range (Fig. 3B; table S9) (ANOVA, host treatment: F(1,822)=24.067, p=<0.001). The degree of population divergence, measured as the sum of allele frequency change (ANOVA, host treatment: F(1,28)=12.937, p=0.001); Euclidean genetic distance from the ancestor (ANOVA, F_1,28_ = 8.633, p =0.007; Tukey post-hoc test t = 2.938, p = 0.007); and within-treatment diversification (t=-8.982, p=<0.001) were also reduced across host treatments (fig. S12). We investigated the extent to which the same *de novo* mutations evolved independently across replicates in response to host or temperature-mediated selection, indicative of adaptive evolution (*55*). Evolving within hosts drove the most molecular parallelism (>=2 replicates; Fig. 3C; table S9) (host/environment pairwise t-test=-3.41 p=<0.001), with evidence for variants in genes linked to infection repeatedly arising, persisting, and increasing in frequency across replicates, consistent with selection (Fig. 3D). Higher levels of genetic parallelism are likely to arise when the adaptive mutational target size is low (*56*), such as in extreme environments (*57*) and in parasites under strong host-mediated selection (*58, 59*). Here, our findings are consistent with host-mediated selection constraining the size of the adaptive landscape, dominating population genomic changes relative to temperature.

The impact of temperature on the genetic architecture of adaptation was most distinct in the environmental populations. Warm temperatures facilitated the fastest rates of evolution, with the largest changes in mean allele frequencies genome-wide (Fig. 3B; table S9). However, these evolutionary rates do not align with replication rates across temperatures (*60*) – both *in vitro* and within-host levels of evolved replication were highest at 30°C (figs. S2, S5, S13) (one-way ANOVA, 25°C/30°C, t=4.14 p=0.0002). Environmental populations evolving in warm conditions also showed relaxed evolutionary constraint, with higher rates of allele frequency change for non-parallel (unique to a single lineage) mutations (Fig. 3C) (linear model parameter estimate, β=-0.045, DF=822, t=-2.753, p=0.006). In contrast, parasite populations in cool and heatwave conditions experienced the lowest average allele frequencies (Fig. 3B) and nucleotide diversity (ANOVA, heatwave parameter estimate, t=-2.681, p=0.012; fig. S14). Analogous to all host-adapted populations, SNPs that did spread at extreme temperatures in the environment were more often parallel across replicate populations, consistent with a subset of alleles with strong phenotypic effect size involved in convergent adaptation (Fig. 3C). Our genomic results suggest large effect responses at few loci and polygenic evolution may occur to differing extents in response to abiotic and biotic pressures. While functional validation of specific candidate genes identified here may contribute to genomic surveillance efforts during climate-driven epidemics, polygenicity in pathogen evolution may pose a challenge to these efforts (*61*). Nonetheless, our results suggest pathogen evolution may be more predictable, and thus manageable, when monitoring infectious diseases with climate change.

Our study reveals that temperature can drastically shape the virulence and genetic architecture of adaptation for bacterial parasites, whether in a host or the environment. We found that warm temperatures typical of the natural environment for the tropical parasite maximized the capacity for a virulence boost once in a new temperate host. The host-parasite thermal mismatch further imposed selection on parasites with constant heat favouring or maintaining virulent variants that are only revealed at cooler temperatures or when host tolerance subsides. The increasing severity of temperatures and novel infection are already placing many species at risk (*24, 37, 62*). The rapid evolution of potentially cryptic virulence will intensify these threats. Yet, any reduction in the pace of adaptation at more extreme temperatures could increase the window of opportunity for the development of effective clinical interventions to emerging infectious diseases. Further investigations should assess whether the timing of mass mortality events in animal populations is similarly linked with extreme temperature patterns. Going forward, we should consider climate data alongside geographic range overlap (*63*), host phylogenetic relatedness (*64, 65*), and environmental residency (*33*) in epidemiological (*7*) and evolutionary predictions of outbreaks. This focus could be particularly informative for getting ahead of virulent variants emerging at the edge of expanding ranges or within existing host habitats where temperatures are becoming extreme (*4*–*6, 66*).

## Acknowledgments

We are grateful to Kieran Bates and Jonathan Hodgkin for advice on *Leucobacter musarum*. We thank members of the King lab for insightful discussions.

## Funding

Philip Leverhulme Prize (KCK)

NERC Environmental Omics Facility grant (no. NEOF1368) (KCK, TEH)

NERC Research Project Grant (NE/X000540/1) (KCK)

NSERC Canada Excellence Research Chair (KCK)

## Author contributions

Conceptualization: TEH, KCK

Investigation: TEH, JCF, JL, KLH, EJS, SJ

Data curation: TEH, JMK, JF, EJS

Formal analysis: TEH, JMK, KLH

Funding acquisition: KCK, TEH

Writing – original draft: TEH, KCK

Writing – review & editing: TEH, KCK, JMK, JCF, JL, KLH, EJS, SJ

## Data and materials availability

Raw sequences deposited in the NCBI Sequence Read Archive (SRA) under BioProject: PRJNA1208452, and processed data will be deposited in the Dryad Digital Repository.

## Supplementary Materials

Materials and Methods

Supplementary Text

Figs. S1 to S14

Tables S1 to S9

References (67–87)

Appendix 1: Detailed experimental protocols

