## Supplementary material for "Rising temperatures favour parasite virulence and parallel molecular evolution following a host jump": Supplimentary file

**The PDF file includes:**

Materials and Methods
Figs. S1 to S14
Tables S1 to S9
References (67–87)
Appendix 1: Detailed experimental protocols

**Other Supplementary Materials for this manuscript include the following:**

Movies S1 to S#
Audio S1 to S#
Data S1 to S#

### Materials and Methods

Host-parasite system

We used a natural bacterial parasite of Caenorhabditis nematodes, Leucobacter musarum sp. nov. subsp. musarum subsp. nov. strain CBX152T (35). Leucobacter musarum is a gram-positive bacterium isolated from infected C. tropicalis nematodes (JU1635 strain), collected from rotting banana trunks on Cape Verde (Republic of Cabo Verde) ((45), Main text Fig 1).

In this study we used simultaneous hermaphroditic N2 wild-type C. elegans strain from the Caenorhabditis Genetics Centre (University of Minnesota, Minneapolis, MN).

Preparation of host population stocks and ancestral bacteria

Nematode maintenance followed standard protocols ((67); See Appendix for detailed protocols for each procedure mentioned below). Prior to our evolution experiment, a population of C. elegans (N2 strain) was defrosted from glycerol stocks and grown on nematode growth medium (NGM) in 9 cm petri plates seeded with Escherichia coli OP50 (Caenorhabditis Genetics Centre, University of Minnesota, Minneapolis, MN). This food bacterium was grown at 30°C shaking at 200 rpm overnight in LB broth, with 100µl of culture subsequently spread onto each NGM plate and incubated overnight at 30°C. Plates of C. elegans containing gravid females were "bleached" using a sodium hypochlorite solution to surface sterilize eggs and synchronize the population (67). Aliquots of hatched L1 worms were subsequently frozen in sterile worm freezing medium at -80°C. These frozen worm stocks were used throughout the evolution experiment as an evolutionary static host population.

To obtain an ancestral Leucobacter musarum clone, bacteria were streaked from -80°C stocks frozen in 50% glycerol onto LB agar in a 9cm petri dish. The plate was incubated at 25°C for 48 hours until district colonies were visible. A single colony was picked into 8 ml LB broth in a 15 ml falcon tube and grown overnight at 25°C shaking at 150 rpm. This overnight culture represented our ancestral L. musarum. A portion was frozen at -80°C in 50% glycerol, and the remainder used to commence the first passage of the evolution experiment.

Experimental evolution

We passaged L. musarum under four temperature treatments (20°C, 25°C, 30°C, and 25°C-35°C heat shock) and two host treatments (hosts present and environmental/no-host controls) (see Main Text Fig 2). For each of the four host treatments, we independently passaged six biological replicate lines (24 lines total), and three replicate populations per treatment for the environmental controls (n=12). Except for the treatment temperatures during experimental exposures, all in vitro culturing of L. musarum was done at a common temperature (25°C – ambient for L. musarum), E. coli OP50 at 30°C, and all nematode maintenance at 20°C (optimal for C. elegans), to minimise and standardise laboratory selection.

For the experimental exposure plates, we standardised the ancestral L. musarum culture to an OD630 of 0.2, and an overnight culture of E. coli to OD630 of 0.4. The two bacterial cultures were then combined with one-part L. musarum and four parts E. coli (1:4), and 200ul was spread on 90mm NGM plates (host present treatments), or 200ul of L. musarum alone for environmental no-host control treatments. All plates were incubated overnight at 25°C.

Prior to experimental infections, recently defrosted nematodes from our standardised population were surface-sterilised, and age-synchronized. Eggs were allowed to hatch overnight in M9 media at 20°C. After hatching, L1 larvae were spotted onto lawns of E. coli grown on NGM plates (approx. 2500 L1s per plate) and allowed to develop to young adults at 20°C (approx. 60 hours). For experimental parasite exposures 1000 young adult worms were spotted onto each "host present" experimental exposure plate and allowed to dry under sterile conditions. All plates were sealed with Parafilm and placed at their respective treatment temperatures for 30 hours. All no-host control lineages were treated identically except for the addition of nematodes.

After the 30-hour infection period, all plates were taken out of their respective treatment temperatures and kept at room temperature. For each "host present" lineage, 60 worms were picked randomly with a sterile platinum wire into 1.5ml Eppendorf tubes containing five to ten 0.5mm zircon beads and filled with 1200ul M9-Tx. Once all lines had been picked, worms were centrifuged at 1000 rpm for one minute, 1000ul of the supernatant was removed, and 1000ul sterile M9-Tx was added. This process was repeated two more times to gently wash worms of excess surface and lawn bacteria. After the final wash step (using M9 without Triton-X), worms were disintegrated in a tissue lyser in 200ul of M9. Lysed worm solutions were then diluted to 10-4 in sterile M9, and 100ul of this dilution was spread onto a 9cm LB agar plate using glass beads. For no-host control linages, bacteria were transferred directly onto 9cm LB petri dishes from the experimental exposure plates using inoculation loops. All plates were subsequently incubated at 25°C for 48 hours.

Once colonies had grown an inoculation pick was used to pick 100 colonies (host present treatments) or a streak taken (no-host controls) from each lineage into 3 ml LB broth in 15 ml falcons. Tubes were incubated overnight at 25°C shaking at 150rpm. A portion of each overnight culture was frozen in 50% glycerol at -80°C, and the remainder used to make exposure plates for the next passage.

Host survival and parasite CFU assays

To assess changes in parasite infection traits, we performed phenotypic assays using lineages from passage 10 alongside the ancestral culture.

Age-synchronized young adult nematodes were prepared as described above. Bacterial parasite lineages were inoculated directly from frozen stocks into LB broth and grown for 18 hours at 25°C. Exposure plates were prepared by as described above. For assays we used either 60mm or 35mm NGM plates. Approximately 100-200 young adult nematodes were transferred to each exposure plate and then incubated at the appropriate assay temperature. After ~36 hours, the number of live and dead nematodes was counted using a dissecting microscope. Worms were scored as dead if they did not move in response to being prodded by a platinum wire.

Colony forming units (CFU), a proxy for infection burden, were measured following a similar protocol to that used during experimental evolution (see above and Appendix). After 36-hour experimental infections, 10 worms per replicate were randomly picked using a sterile platinum wire into 1.5ml Eppendorf tubes containing zircon beads and M9-Tx. Worms were washed three times before being crushed with a tissue lyser. Samples were serial diluted to 10-4 and various dilutions were spotted onto LB agar plates and incubated at 25°C for 48-hours.

Colonies were then counted and used to calculate the average number of L. musarum bacteria per worm.

In vitro growth assays

Growth curves were obtained by measuring the optical density (OD). Ancestral and evolved cells were initially cultured from frozen populations into LB broth at 25°C overnight. Subsequently, the OD600 of the bacterial samples were standardized and transferred into wells of a transparent 96-well plate. The optical density was recorded using a BioTek Synergy H1 microplate spectrophotometer (Agilent, Santa Clara, CA). Between the OD measurements, which occurred approximately every 10 minutes, orbital shaking was performed to ensure uniform cell distribution. Growth metrics were mathematically computed using the 'gcplyr' R package (68).

Biofilm assays

Biofilm formation was assessed using a crystal violet staining method adapted for 96-well microtiter plates. Bacterial cultures were grown in 96-well plates with a total volume of 200 uL per well. The plates were incubated under the desired experimental conditions for 24 hours. After incubation, the culture supernatants were carefully removed and then gently washed twice with 200 uL of phosphate-buffered saline (PBS, pH 7.4) to remove non-adherent cells. To stain the biofilms, 100 uL of 0.1% (w/v) crystal violet solution was added to each well. The plates were incubated at room temperature for 30 minutes to allow the dye to bind to the biofilm matrix. Following staining, the crystal violet solution was carefully removed by pipetting, and the wells were washed twice with PBS to eliminate excess unbound stain. The bound crystal violet was resolubilized by adding 100 uL of 33% (v/v) acetic acid to each well. The plates were then placed on a shaker and agitated gently at room temperature for 20 minutes to ensure complete dissolution of the dye from the biofilm matrix into the solution. The optical density (OD) of the solubilized crystal violet in each well was measured at a wavelength of 575 nm using a microplate reader. The absorbance values correspond to the amount of biofilm formed by the bacterial cultures under the tested conditions.

DNA extraction and sequencing

For pooled samples, we grew 40 individual colonies for each replicate population separately at 25˚C for two days, then standardized the OD600 of each individual colony before pooling them into one tube to perform DNA extraction. We extracted genomic DNA using a DNeasy Blood and Tissue Kit (Qiagen). Briefly, we pelleted 1ml of each bacterial sample, removed the supernatant, then resuspended in 180ul of enzymatic lysis buffer (20 mM Tris-HCl, 2 mM EDTA, 1.2% Triton, and 3.6mg lysozyme). Samples were incubated at 56˚C for 30 minutes with occasional vortexing. We added 25ul Proteinase K and 200ul Buffer AL to each sample, then incubated them at 56˚C for 30 minutes to one hour. After vortexing for 15 seconds, we added 200ul of 100% ethanol and vortexed briefly before transferring to spin columns. We then followed the manufacturer's instructions for washing and eluting DNA. We quantified DNA concentration using a Qubit fluorometer. We assessed DNA purity and integrity using a Nanodrop and gel electrophoresis, respectively.

Ancestral assembly and variant calling

Ancestral assembly and variant detection were carried out by Xuan Liu at the University of Liverpool Centre for Genomic Research. Initial processing and quality assessment of the sequence data was performed using an in-house pipeline developed by Dr Richard Gregory. Briefly, base calling and de-multiplexing of indexed reads was performed by CASAVA version 1.8.2 (Illumina) to produce 131 samples in FASTQ format. The raw FASTQ files were trimmed to remove Illumina adapter sequences using Cutadapt version 1.2.1 (69). The option "-O 3" was set, so the 3' end of any reads which matched the adapter sequence over at least 3 bp was trimmed off. The reads were further trimmed to remove low quality bases, using Sickle version 1.200 with a minimum window quality score of 20. After trimming, reads shorter than 20 bp were removed. Trimmed reads files were deposited in the NCBI Sequence Read Archive (SRA) under BioProject: PRJNA1208452 (http://www.ncbi.nlm.nih.gov/bioproject/1208452).

The ancestral isolate was assembled using SPAdes (70) and was assessed using BUSCO (71). The assembly was further annotated using Prokka (72).

R1/R2 reads from each sample were mapped to the ancestral assembly using BWA mem version 0.7.5a (73) with default parameters. To retain only confidently aligned reads, alignments were filtered to remove reads with a mapping quality lower than 10, which equates to a 10% chance that the read was derived from another genomic location. Duplicate reads arising from PCR amplification can bias variant calling. To avoid this, read duplicates were identified and filtered to retain only a single representative, using the Picard "MarkDuplicates" tool, version 1.85 (http://picard.sourceforge.net/).

In order to quantify putative functional effects of de novo mutations throughout our evolution experiment, we initially called SNPs using the Freebayes (74) using default settings except ploidy settings. "ploidy= 1" was set for clonal samples, whilst "ploidy = 40" was set for population samples. A "QUAL > 20" filter was also applied. We then annotated variants using SnpEFF (version 4.3) (75).

Population genomic analysis

To quantifying genomic responses over the course of the evolution experiment, we called the frequency of de novo mutations in sequenced reads at the experimental endpoint. Note that we are able to ascribe all SNPs as de novo mutations, as the reference genome to which all evolved lines were aligned to was generated from the ancestor used to start the experiment. As sequence reads represent pools of clonal populations in each experimental and control replicate, we leveraged a highly efficient recoding of the popular pipeline for experimental evolution projects to do so (PoPoolation2: (76)) called Grenedalf (77). We used the --frequency command to produce a table of de novo mutation frequencies at each site in the genome from bam alignment files. As these mutations were absent at the beginning of the experiment, the frequency of these mutations at the end of the experiment also represent their allele frequency change. We also used the Popoolation2 implementation to calculate diversity calculated as the pairwise diversity at each locus. As each host-evolved treatment had 6 experimental replicates for each of the 4 temperature regimes, and each no-host (control) treatments had 3 experimental replicates for each of the 4 temperature regimes, we attained allele frequency and diversity estimates at each locus in 36 distinct pools.

To quantify how temperature and host regimes facilitate or constrain rates of pathogen evolution, we calculated genomic divergence from the ancestor in each experimental replicate and treatment, by summing the frequency of allele frequency change across all loci (Figure S13A). Additionally, we calculated the a matrix of Euclidian distances across treatments for allele frequency change at all loci. Euclidian from the ancestor was calculated as the distance from zero for each loci, alongside each within-treatment pairwise distance (Figure S13B and C).

As we aimed to investigated signals of adaptive evolution over the course of the experiment, we took two approaches. One approach was to identify mutations present across multiple replicates given host status, temperature status, and host x temperature status to identify those involved in convergent, repeatable genomic evolutionary response. The second approach was to identify de novo mutations that have dramatically increased in the frequency over the course of the experiment. For the 810 de novo mutations present across at least one replicate, we quantified the number of replicates it was present in, which temperature regime it evolved under, which host regime it evolved under, and its location in the genome relative to the reference. For all following analyses, we combined these approaches, by only including variants that either displayed significant repeatable evolution within an experimental treatment, or failing that, had minor allele frequencies (MAF) of at least 10%. Doing so meant we could focus on loci most likely to be under selection, without omitting potentially important rare mutations.

To test whether genome-wide allele frequency change and diversity (in the absence of recombination and migration, due to only mutation, selection, and drift) corresponded to experimental treatment, we performed a multiple linear regression to test whether host, temperature, and host x temperature, and presence across experimental replicates explained each independent variable. We expected that mutations that appeared across multiple replicates would be those most likely representing a convergent response to selection. If so, we might expect that such mutations also show the most allele frequency change over the course of the experiment. We performed these multiple regression analyses using the lm function and a type 3 Anova in R (78). As the number of experimental replicates varied across host and non-host treatments, in both models we normalized replicate number as a predictor such that it would be comparable across treatments. We then extracted the least squares means from the multiple regression analysis, using the function lsmip in R. (79) This allowed us to pull out the mean temperature and host effect on allele frequency change/diversity, while accounting for the modelled interaction effect.

We also tested whether particular loci show significant parallel responses to specific temperature regimes when evolved in the host context (Fig S10). To do so, we leveraged the six independent experimental replicates for each temperature treatment to quantify the mean and standard error of allele frequency change, and calculated the corresponding test statistic and significance based on a p-value cutoff of alpha=0.05.

Statistical analysis of phenotypic data

All statistical analyses were performed in R [v. 4.2.1 (80)]. Data wrangling and figures were produced using the "Tidyverse" (81), "cowplot" (82), "ggpubr" (83). Descriptions of model structure and error distributions for statistical models are shows below (Table SM1). Significance of fixed effects were assessed with Analysis of Variance (ANOVA, Type III; 'car' package (78)). Post hoc pairwise tests were performed using the 'emmeans' package (79)

To analyse the relationship between virulence and within-host burden (Main Text Fig 2C) we first predicted average trait values for host mortality and CFUs (predictions using models a, e, f, & g in Table SM2). For host-evolved lines, we predicted for each level of the random effect, that is, we extracted the mean predicted values for each replicate line. For environment-evolved treatments, we predicted the mean treatment-level main effect. In this analysis we were interested in evolutionary change that is specific to association with a host. We therefore calculated the relative host effect of both traits for each host lineage by subtracting the predicted mean of their respective no-host control. For each assay temperature both traits were then centered and scaled prior to analysis with Multivariate Analysis of Variance (MANOVA; 'car' package (78)). Pearson's correlation was used to assess the within-treatment correlation.

**Table SM1 (separate file).**

Details of statistical models analysing host mortality or colony forming units (CFU) phenotype data. Each model assessed the impact of experimental "Evolution temperature" treatment, "Host treatment", "Assay temperature" (temperature at which traits were measured at the end of the experimental evolution), or combinations thereof, as fixed effects. Generalized Linear Mixed Models were fitted using the "glmmTMB" package (84), and Linear Mixed Models using "lme4" package (85). Fixed and random effect structures were determined by experimental design (maximal models) and model fit diagnostics were assessed using the "DHARMa" package (87). Temperatures include 20°C, 25°C, 30°C, and the experimental heatwave (HW, see methods). Model structures are shown using R syntax (additive effect = "+", interaction = "*", nested random effect = "/"). Where appropriate the significance of fixed effects were tested using Analysis of Variance (ANOVA type III) using the "car" package (78).

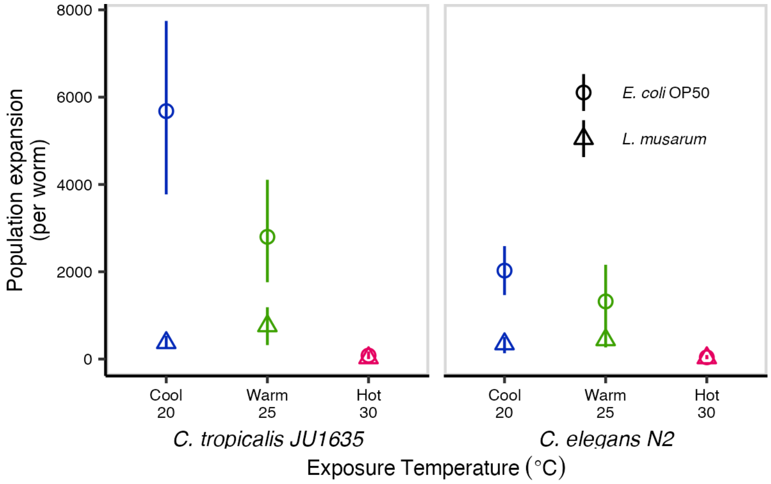


**Fig. S1.**

Lineage expansion/population size of *C. elegans* (N2 strain) and *C. tropicalis* (strain JU1653) worms across two generations following exposure to *L. musarum* at 20°C, 25°C, or 30°C. After 24 hours exposure to the parasite or food (*E. coli* OP50), five worms per replicate were picked onto fresh plates containing just food. Each worm population was then allowed to expand for 3-4 days before assessing population size (at 20°C for *C elegans* and 25°C for *C. tropicalis*).

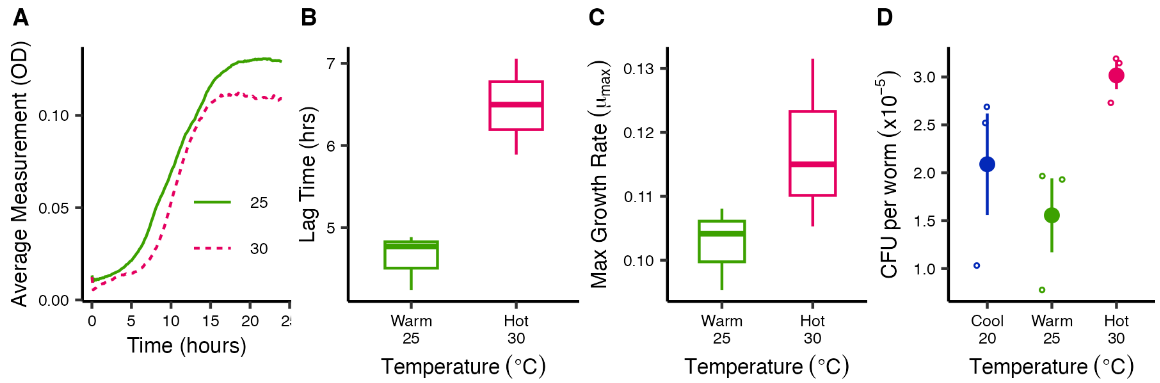


**Fig. S2.**

*In vitro* growth metrics (OD630) and within-host burden for the ancestral parasite clone across temperature. A) Average optical density ($OD_{600}$) measurements across time for *L. musarum* samples grown at either 25 or 30°C. B) Average lag time at each temperature. C) Average maximum growth rate. Metrics were calculated using the gcplyr R package (*68*) (see methods). D) Colony Forming Units (CFU) – a measure of within-host bacterial load – of ancestral parasite isolate across three constant environmental temperatures. While the effect of temperature here was marginally non-significant (Kruskal-Wallis Test: statistic = 5.956, d.f. = 2, p-value = 0.051), there was a trend towards increased numbers of within-host bacterial cells at 30°C (Dunn Test 25°C-30°C: statistic = 2.385, p-value = 0.017, adj. p-value = 0.0512).

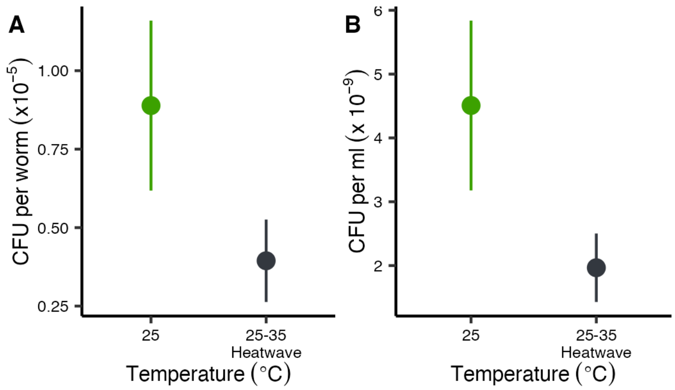


**Fig. S3.**

Viability of *Leucobacter musarum* following heat shock. A) *In vivo* cell viability. Colony forming units within host *C. elegans* after exposures at either 25°C or 35°C heat shock (see Supp. Materials). The trend for reduced CFU following heat shock was not significant (Kruskal-Wallis Test: statistic = 1.191, d.f. = 1, p-value = 0.275). B) *In vitro* cell viability. Twelve overnight culture of *L. musarum* grown from individual colonies were distributed evenly across three 96-well plates and maintained at 25°C or exposed to a three-hour heat shock at 35°C. Following the heat shock, all replicates were spotted on LB agar and incubated at 25°C for 48 hours, after which the number of colonies were counted. There were significant fewer viable cells following heat shock (Kruskal-Wallis Test: statistic = 14.552, d.f. = 1, p-value = <0.001).

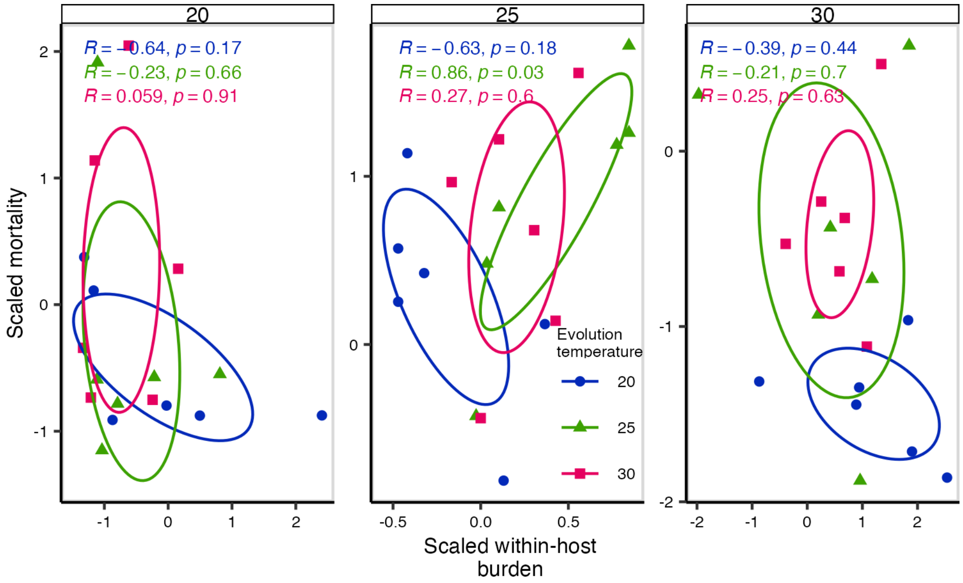


**Fig. S4.**

Raw data (scaled, and centered on ancestor) for colony formatting units (within-host burden) and virulence (mortality) of *L. musarum* evolved within hosts at 20, 25, or 30°C. Traits were measured for populations from all treatments at each temperature in subsequent assays (facets). Coloured text shows Pearson's correlation coefficients for each treatment and temperature (20°C = blue, 25°C = green, 30°C = pink). As with the data shown in Main Figure 2C, there is only a positive relationship between within-host burden and host mortality for parasite populations passaged at warm 25°C.

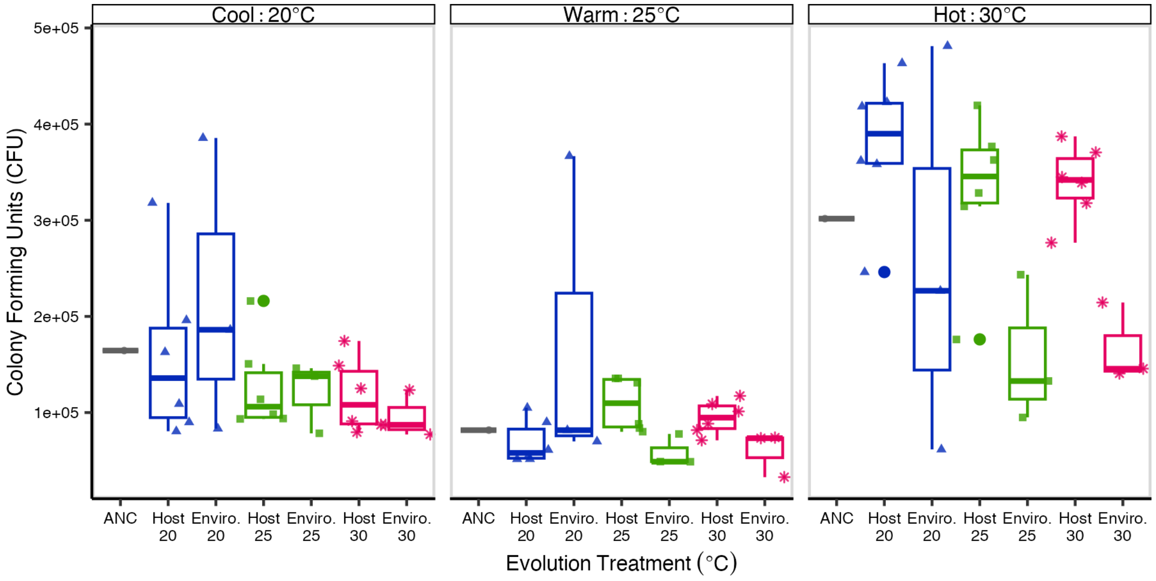


**Fig. S5.**

Within-host colony forming units (CFUs) depend on evolution treatment and assay temperature. A) within-host CFUs from nematode hosts exposed to the ancestral isolate (ANC) or parasite populations passaged either within hosts (host) or environmental controls (enviro) at either 20, 25, or 30°C. Parasite CFUs of all populations were assayed as each constant temperature (facets).

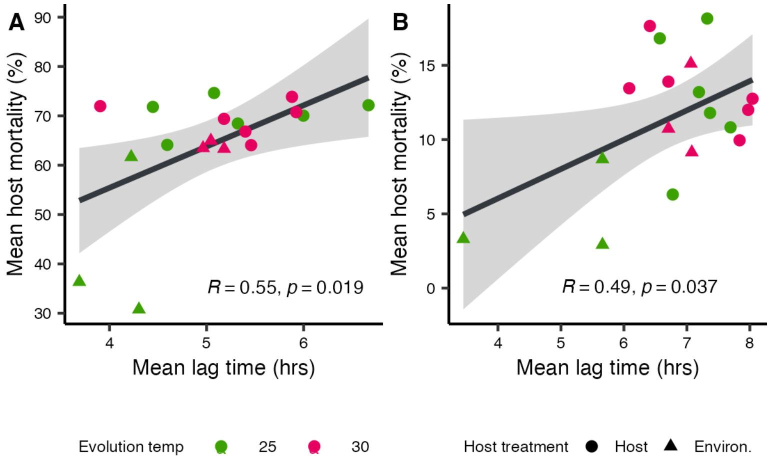


**Fig. S6.**
The relationship between mean in vitro lag time and mean virulence (host mortality) for each population passaged at 25°C (green) and 30°C (pink) within-hosts (circles) or environmental no-host controls (triangles), at A) 25°C, or B) 30°C assay temperatures. Inset text shows Pearson's correlation coefficients.

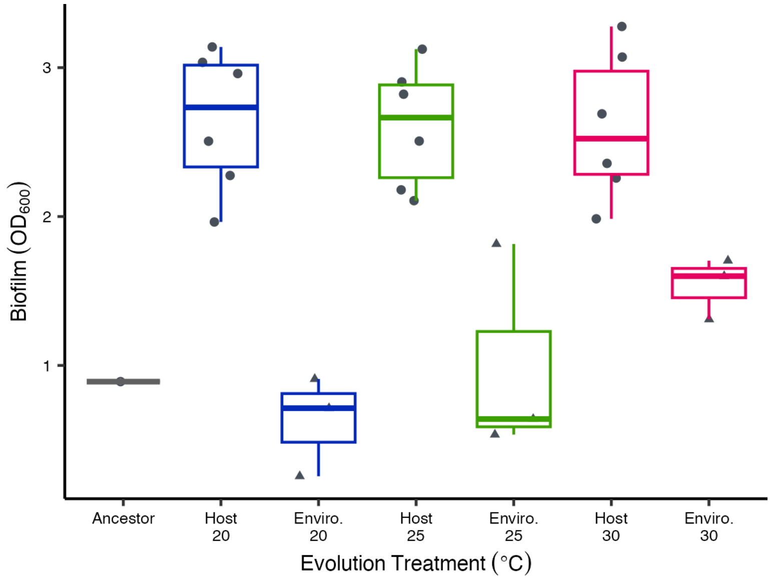


**Fig. S7.**

Increased biofilm formation following within-host evolution. Average biofilm formation (measured as optical density 600nm) for evolved parasite populations and the ancestral isolate. Biofilm formation was measured for populations grown at 25°C. Host = within-host treatments, Environ. = environmental controls, ANC = ancestral isolate.

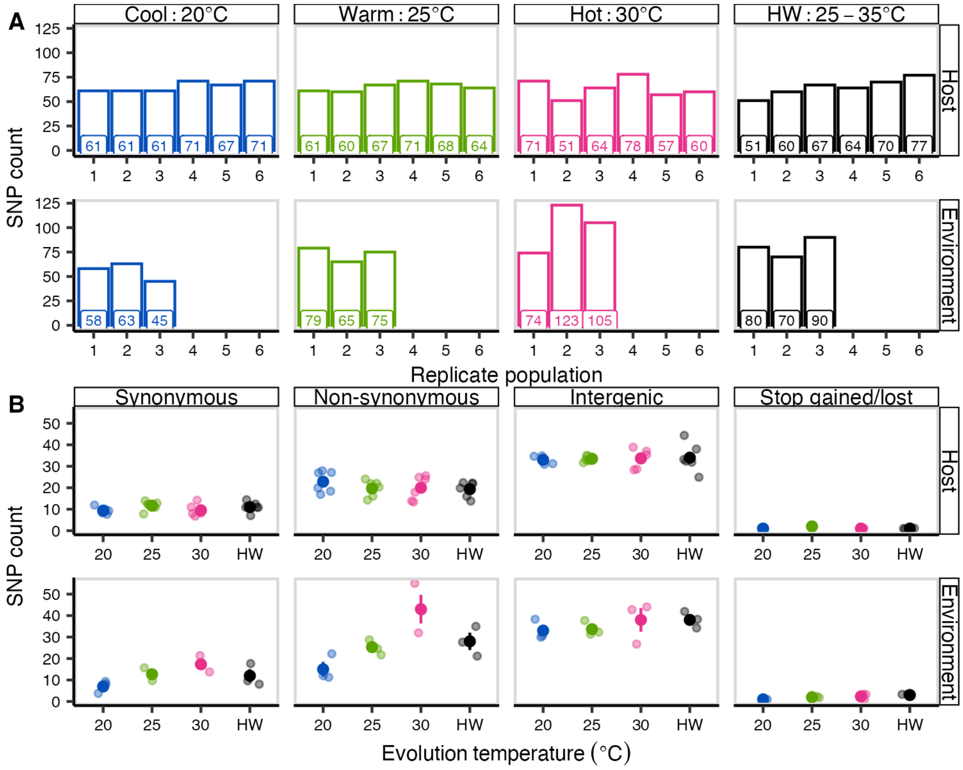


**Fig. S8.**

Single nucleotide polymorphism (SNP) counts for evolved parasite populations from each treatment. A) Counts of variants (genome-wide) for each evolved lineage from pools of 40 sequenced clones. The x-axis shows each independent population. B) Average SNP count (±SE) per treatment for different mutation type. Transparent points show counts for each independent population.

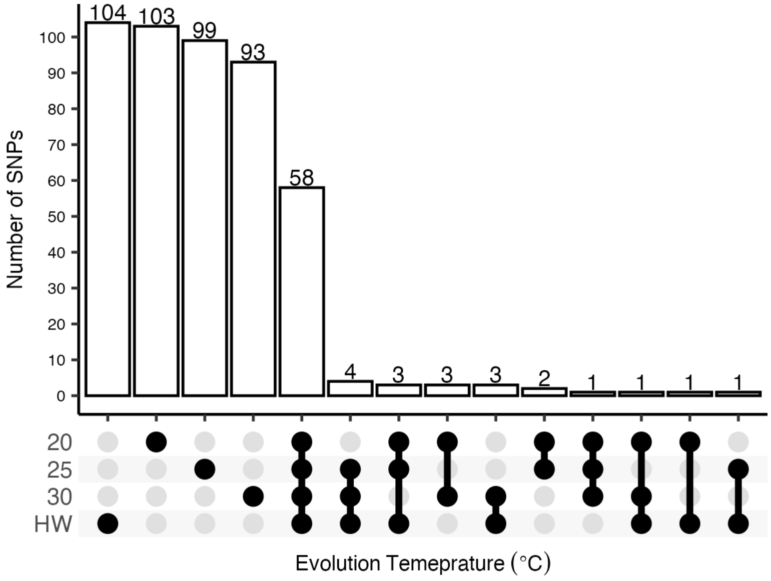


**Fig. S9.**

Most single nucleotide polymorphisms (SNPs) are either shared across all treatments or unique to a single treatment. Shown are the number of SNPs (genome-wide) that occurred in within and across different combinations of treatments for host-evolved parasite populations.

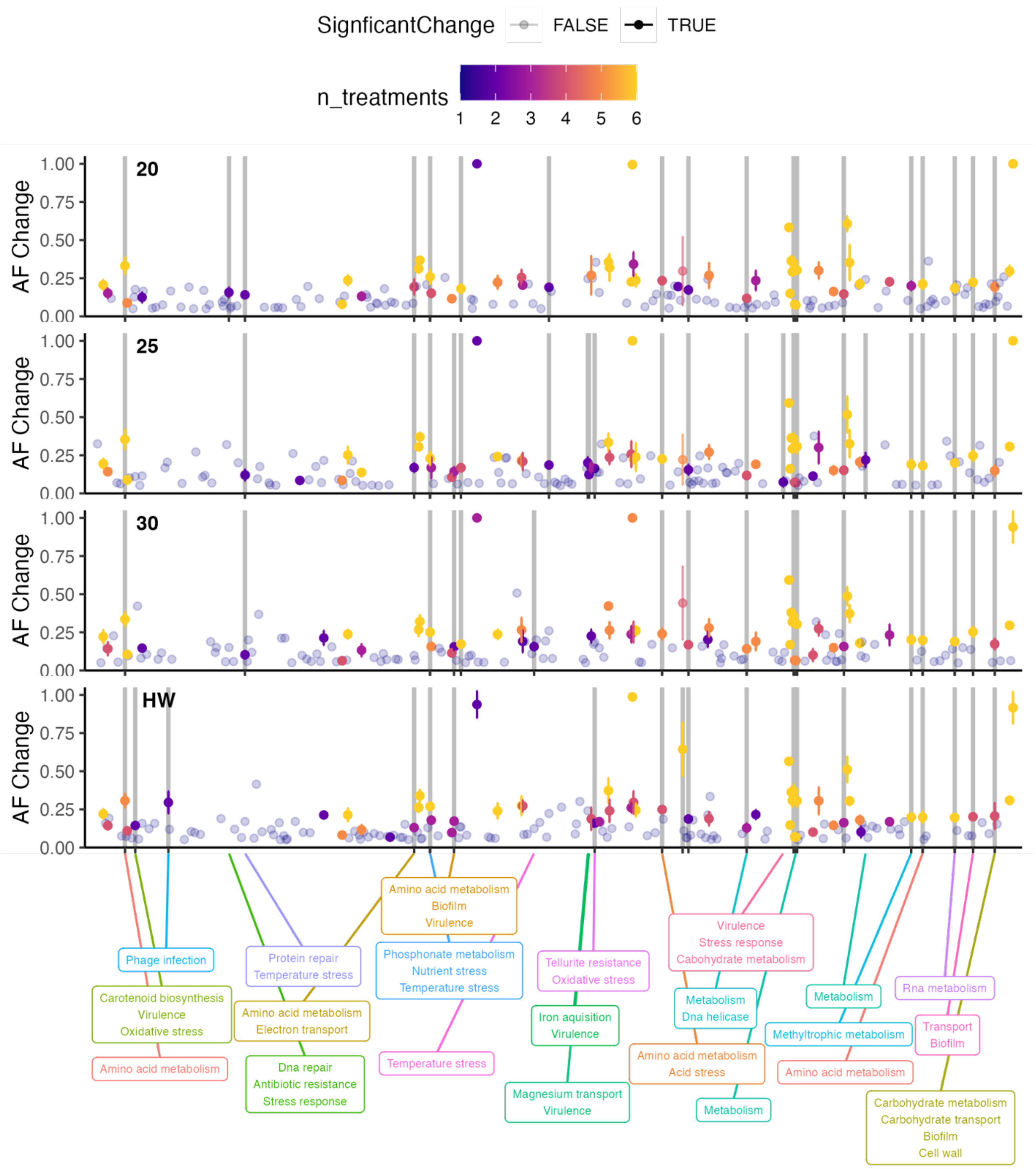


**Fig. S10.**

Mean allele frequency change across host evolved populations from each evolution temperature. Shown are SNPs along the genome. Colour represents the number of populations a SNP arose in parallel, non-translucent points showed significantly higher allele frequencies than would be expected at random in parallel SNPs. Grey bars show the position of non-synonymous genic SNPs which showed significant allele frequency change in parallel across multiple populations (with the corresponding putative functions below).

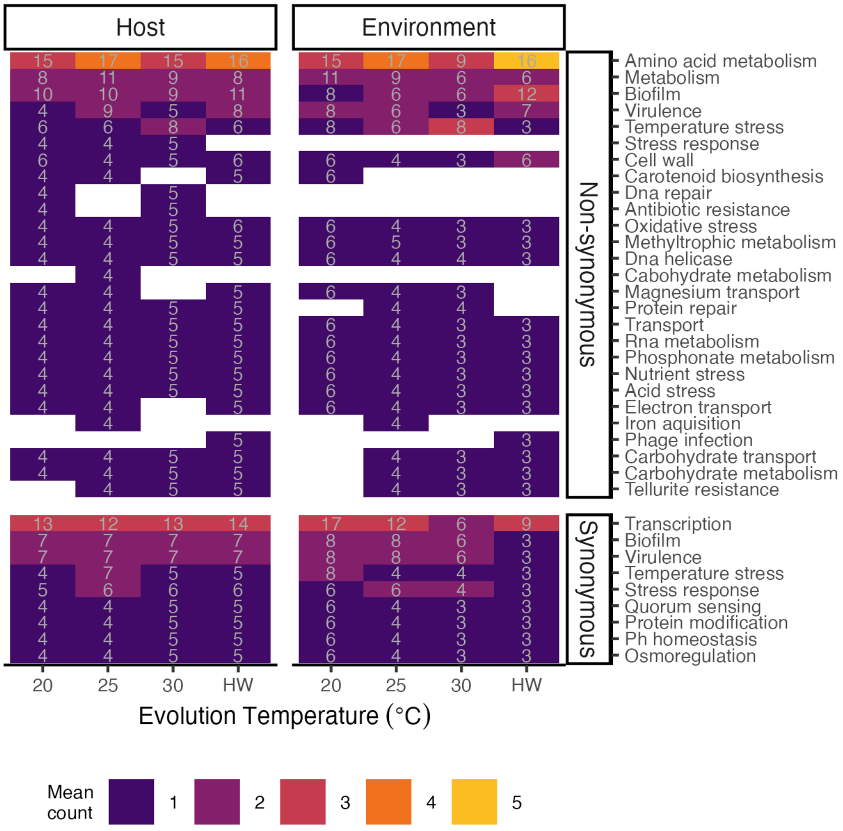


**Fig. S11.**

Putative functions and number of genes potentially under selection. For each evolution treatment, shown are the average SNP counts (colours) and average proportion relative to all SNPs in that treatment (numbers) for each putative functional category. Only SNPs within genes are shown. The data was further filtered to only include SNPs with minor allele frequencies >10% or variants which showed significant parallel evolution across replicate populations within treatment.

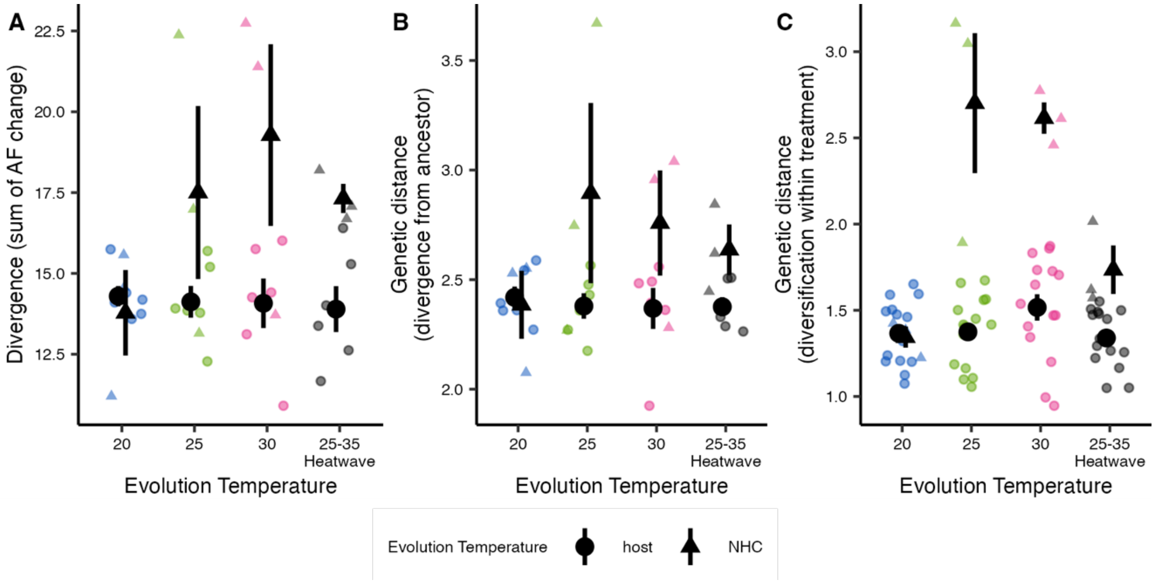


**Fig. S12.**

Genetic distance and divergence of ‘host-evolved parasite populations across temperatures. A) Genetic divergence as the sum of allele frequency change for populations in each treatment, B) Euclidean distance of each evolved population to the ancestor from all SNPs (MAF >0.1). C) Euclidean distance between populations within each evolution treatment, showing within treatment diversification.

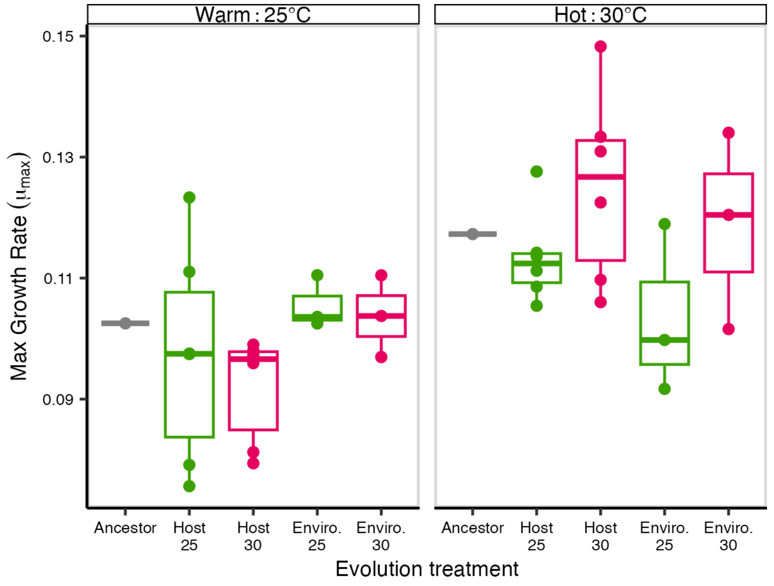


**Fig. S13.**

*In vitro* growth rates for parasite populations at two test temperatures (25°C & 30°C). Shown are the calculated maximum growth rates ($\mu_{max}$) for evolved parasite populations or the ancestral isolate during *in vitro* growth in 96-well plates at either 25°C or 30°C (facets). Both 'host-evolved' (Host) and environmental controls (Environ.) are shown.

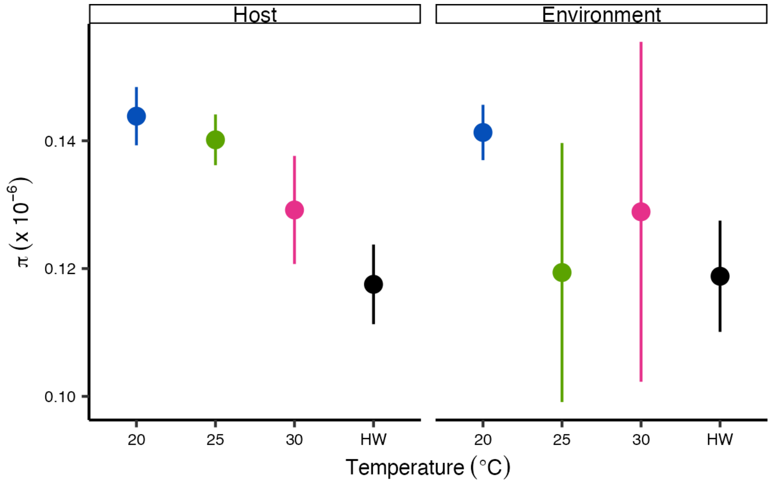


**Fig. S14.**

Nucleotide diversity for 'host-evolved' parasite populations declines at the hottest temperature treatments. Per gene nucleotide diversity, predicted using the PoPoolation2 software (*76*) following the standard workflow (https://sourceforge.net/p/popoolation2/wiki/Main/).

**Table S1.**

Virulence of constant temperature evolved lines relative to the ancestor. Significance of fixed effects (Analysis of Variance, type III) for linear mixed effect model for the effects of evolution temperature (20, 25, and 30°C) and assay temperature (20, 25, and 30°C) on mortality for host evolved parasite populations relative to the average ancestral value.

| Term | Chisq | df | p-value |
|---|---|---|---|
| **Evo temp** | 12.4997 | 3 | 0.006 |
| **Assay temp** | 10.3142 | 2 | 0.006 |
| **Evo temp:Assay temp** | 2.3159 | 4 | 0.678 |

**Table S2.**

Parameter estimates from a linear mixed effect model testing the effects of evolution temperature (20, 25, and 30°C) and assay temperature (20, 25, and 30°C) on mortality for host evolved parasite populations relative to the average ancestral value (ancestor average set as intercept).

| Assay temp | Evo temp | Estimate | s.e. | d.f. | t-value | p-value |
|---|---|---|---|---|---|---|
| **20** | 20 | -2.366 | 1.532 | 45 | -1.544 | 0.130 |
| | 25 | -1.379 | 1.532 | 45 | -0.900 | 0.373 |
| | 30 | 1.306 | 1.532 | 45 | 0.853 | 0.398 |
| **25** | 20 | 1.352 | 1.532 | 45 | 0.882 | 0.382 |
| | 25 | 4.054 | 1.532 | 45 | 2.646 | 0.011 |
| | 30 | 3.328 | 1.532 | 45 | 2.173 | 0.035 |
| **30** | 20 | -6.881 | 1.532 | 45 | -4.491 | <0.001 |
| | 25 | -2.428 | 1.532 | 45 | -1.585 | 0.120 |
| | 30 | -1.989 | 1.532 | 45 | -1.298 | 0.201 |

**Table S3.**

Virulence (host mortality) of parasite populations passaged at constant temperatures during assays across the three constant temperatures (20, 25, and 30°C). Shown are the significance of fixed effect estimated from a model of host mortality, predicted by evolution temperature treatment (levels = "20", "25", and "30°C"), host evolution treatment (levels = "within-host" and "environmental control"), assay temperature (levels = "20", "25", and "30°C"), and all higher order interactions. The data were modelled using a generalized linear mixed effect model (GLMM) with a Beta-binomial (log link) error distribution using the "glmmTMB" package. Fixed effects were estimated with Analysis of variance (ANOVA, type III), using the "car" package.

| Term | Chisq | df | p-value |
|---|---|---|---|
| **(Intercept)** | 75.7583 | 1 | <0.001 |
| **Evo temp** | 35.6966 | 2 | <0.001 |
| **Host treat** | 59.5085 | 1 | <0.001 |
| **Assay temp** | 745.6287 | 2 | <0.001 |
| **Evo temp:Host treat** | 17.9014 | 2 | <0.001 |
| **Evo temp:Assay temp** | 0.7778 | 4 | 0.941 |
| **Host treat:Assay temp** | 2.5262 | 2 | 0.283 |
| **Evo temp:Host treat:Assay temp** | 1.1519 | 4 | 0.886 |

**Table S4.**

Pairwise comparisons between parasites evolved within hosts or environmental controls at each level of evolution temperature and assay temperature. Estimated from the GLMM reported in Table S3.

| Assay temp | Evo temp | Log odds ratio | s.e | z-ratio | p-value |
|---|---|---|---|---|---|
| **20°C** | 20°C | 2.17 | 0.501 | 3.362 | 0.0008 |
| | 25°C | 2.09 | 0.481 | 3.208 | 0.0013 |
| | 30°C | 1.06 | 0.242 | 0.254 | 0.7993 |
| **25°C** | 20°C | 2.73 | 0.625 | 4.373 | <0.0001 |
| | 25°C | 3.13 | 0.719 | 4.976 | <0.0001 |
| | 30°C | 1.32 | 0.304 | 1.220 | 0.223 |
| **30°C** | 20°C | 1.72 | 0.504 | 1.863 | 0.0625 |
| | 25°C | 2.81 | 0.815 | 3.571 | 0.0004 |
| | 30°C | 1.17 | 0.300 | 0.610 | 0.5422 |

**Table S5.**

Significance of fixed effects from a MANOVA testing the effects of evolution temperature (20, 25, 30°C) and assay temperature (20, 25, 30°C) on the relationship between relative host mortality and relative within-host colony forming units (CFU), for parasite populations passaged within hosts. Relative trait values were produced by subtracting the average environmental control's value from the matching evolution temperature treatment. This relative measure allows the partitioning of changes to phenotypic traits that were specific to within-host processes.

| Term | Pillai test statistic | Approx F Value | df | Num d.f. | Den d.f. | p-value |
|---|---|---|---|---|---|---|
| **Evo_temp** | 1.256 | 38.001 | 2 | 4 | 90 | <0.001 |
| **Assay_temp** | 0.00 | 0.000 | 2 | 4 | 90 | 1.000 |
| **Evo_temp:Assay_temp** | 0.549 | 4.256 | 4 | 8 | 90 | <0.001 |

**Table S6.**

Virulence (host mortality) of parasite populations passaged at 25°C, 30°C, or under the heatwave temperatures during simulated heatwave (25°C - 35°C ) assays. Shown are the significance of fixed effect estimated from a model of host mortality, predicted by evolution temperature treatment (levels = "20", "25", and "30°C"), host evolution treatment (levels = "within-host" and "environmental control"), and their interaction. The data were modelled using a generalized linear mixed effect model (GLMM) with a Beta-binomial (log link) error distribution using the "glmmTMB" package. Fixed effects were estimated with Analysis of variance (ANOVA, type III), using the "car" package.

| Term | Chisq | df | p-value |
|---|---|---|---|
| **Evo_temp** | 0.195 | 2 | <0.001 |
| **Host_treat** | 24.034 | 1 | <0.001 |
| **Evo_temp:Host_treat** | 12.856 | 2 | 0.002 |

**Table S7.**

Pairwise comparisons between parasites evolved within hosts or environmental controls at each level of evolution temperature during a heatwave assay. Estimated from the GLMM reported in Table S6.

| Assay temp | Evo temp | Log odds ratio | s.e | z-ratio | p-value |
|---|---|---|---|---|---|
| **Heatwave** | 25°C | 0.428 | 0.0764 | -4.754 | <0.0001 |
| | 30°C | 0.513 | 0.0943 | -3.632 | 0.0003 |
| | Heatwave | 0.998 | 0.1731 | -0.013 | 0.9892 |

**Table S8.**
Counts of genome-wide de novo variants at different levels of minor allele frequency (MAF). Average counts are shown for each experimental evolution treatment, with the standard deviation (SD).

| Host treatment | Evolution temperature | Mean number of SNPS genome wide (SD) | Mean SNPS MAF >0.1 (SD) | Mean SNPS MAF >0.5 (SD) | Mean number of fixed (MAF =1) SNPS (SD) |
|---|---|---|---|---|---|
| **Host** | 20°C | 65.3 (4.97) | 51.3 (3.33) | 4.5 (0.55) | 2.2 (0.75) |
| | 25°C | 65.2 (4.26) | 51.2 (3.97) | 3.8 (0.98) | 2.3 (0.52) |
| | 30°C | 63.5 (9.77) | 50.3 (7.97) | 4.2 (1.33) | 2.4 (0.55) |
| | Heatwave | 64.83 (8.89) | 50.0 (7.72) | 4.8 (1.17) | 1.4 (0.55) |
| **Environmental** | 20°C | 55.33 (9.29) | 45.7 (5.78) | 4.7 (1.53) | 1.3 (0.58) |
| **control** | 25°C | 73.0 (7.21) | 54.3 (5.03) | 8.0 (6.24) | 2.7 (0.58 |
| | 30°C | 100.67 (24.79) | 68.7 (15.28) | 6.3 (2.89) | 2.3 (0.58) |
| | Heatwave | 80.0 (10.0) | 63.0 (2.00) | 4.3 (1.54) | 2.3 (0.58) |

**Table S9.**

ANOVA (Type III) of a fully-parameterised model showing the significance of fixed effects for the effect of evolution temperature (20, 25, 30°C, and Heatwave), host treatment (within-host and environmental controls), and level of parallel evolution (No. replicates) on mean allele frequency of variants within evolution treatments. The number of within-treatment replicate populations each de novo variant arose in (No. replicates/Scaled reps) was scaled to between zero and one, to account for the differing number of replicate populations across treatments.

| Term | Sum sq | df | F-statistic | p.value |
|---|---|---|---|---|
| **No. replicates (scaled)** | 0.874 | 1 | 29.4589 | **<0.001** |
| **Evo temp** | 0.460 | 3 | 5.1685 | **0.002** |
| **Host treat** | 0.714 | 1 | 24.0662 | **<0.001** |
| **Scaled reps:Evo temp** | 0.400 | 3 | 4.4933 | **0.004** |
| **Scaled reps:Host treat** | 0.345 | 1 | 11.6257 | **<0.001** |
| **Evo temp:Host treat** | 0.300 | 3 | 3.3736 | **0.018** |
| **Scaled reps:Evo temp:Host treat** | 0.191 | 3 | 2.1448 | 0.093 |

**Data S1. (separate file)**
Type or paste caption here.

**Appendix: Detailed experimental protocols**

*Leucobacter musarum* infection assay schedule

Day 1:
Chunk nematodes to 9cm OP50 NGM plates
Day 4:
Bleach nematodes
Seed 9cm NGM plates with OP50 (enough for ~2000 worms per plate)
Day 5:
Count L1's, and plate on OP50 NGM
Brew OP50 and parasite from frozen stock
Day 6:
Prepare experimental exposure plates
Day 7:
Expose adult worms on exposure plates
Day 8:
Mortality counts or CFU's
Day 10:
For CFU assay count colonies

Media and chemicals

Nematode Growth Medium - NGM (1L)

3g NaCLl
17g agar
2.5g peptone
975ml $dH_20$
1ml 5mg/ml cholesterol
(dissolved in ethanol)
autoclave and allow to cool (to about 50-60° in water bath or hand holding temp.)
1ml 1M $CaCl_2$
1ml 1M $MgSO_4$
25ml 1M Potassium Phosphate
(1L potassium phosphate, pH 6.0 = 868ml 1M $KH_2PO_4$ + 132ml 1M $K_2HPO_4$)
pour plates using sterile technique.

LB agar media

Follow instructions on bottle
16g LB powder
14g Agar powder
800ml $dH_20$
autoclave and allow to cool (to about 50-60° in water bath or hand holding temp).
pour plates using sterile technique.

LB liquid media

20g LB powder
1L $dH_20$

autoclave and allow to cool

1M $CaCl_2$
14.7g $CaCl_2.2H_20$
100ml $dH_20$
Autoclave

5mg/ml Cholesterol
0.25g cholesterol
50ml ethanol

1M $MgSO_4$
24.65g $MgSO_4$
100ml $dH_20$
autoclave

1M $KH_2PO_4$
136.09g $KH_2PO_4$
1L $dH_20$
autoclave

1M $K_2HPO_4$
87.1g $K_2HPO_4$
500ml $dH_20$
autoclave

M9 (1L)
3g $KH_2PO_4$
6g $Na_2HPO_4$
5g NaCl
1L $dH_20$
autoclave and allow to cool (to less than 60° in water bath or hand holding temp.)
1ml $MgSO_4$
If you want to add Triton-x (to make M9-Tx) for worm work add:
1ml 10% Triton-x (in $dH_20$)

5M NaOH (100ml)
20g NaOH
80ml $dH_20$
20ml Mix (swill) until cool (it is an exothermic reaction). Then add more $dH_20$ up to 100ml

Protocols
All protocols assume sterile technique and should be done in either the biological safety cabinet or under the flame unless otherwise stated.

Freezing worms

1. Chunk worms 3-4+ days before freezing
2. Wash starving worms off plates (dauer L2 worms freeze best) using M9 with/without Triton-x into Falcon tubes (see "washing worms off plates")
3. Bleach worms and allow L1s to hatch overnight (see bleaching below)
4. Add 750ul of Worm Freezing Solution to labelled cryovials.
5. Add 500ul of L1s then invert tubes to mix.
6. Put tubes into a polystyrene box and put into the -80°C freezer (so that they freeze slowly).

Thawing frozen worms

1. Take a cryotube from the -80°C freezer. Let it warm to room temperature gradually.
2. Shake tube and Pour/pipette the contents onto OP50 seeded NGM (Some protocols require worms to be put onto unseeded OP50).
3. Let the liquid dry in the biological safety cabinet.
4. Parafilm the plates to ensure they don't get contaminated and pace them in the 20°C incubator.
5. In about a week you should see plenty of worms and can chunk.

Selecting bacterial clones

1. Streak bacteria onto agar plate (LB if OP50 or Leucobacter) using inoculation loop.
2. Streak as the image below shows: change inoculation loop between streak 1 and 2.
3. Place this plate at 30°C overnight (OP50) or 25°C for 48 hours (*L. musarum*).
4. Pick one colony with an inoculation loop.
5. Swill the picked colony with the inoculation loop into liquid LB media (see "Growing bacterial cultures")

Growing bacterial cultures

1. Use inoculation loops to swab frozen samples from cryotubes or a colony from a LB plate (see "selecting bacterial clones") into LB broth. Use the biosafety hood and sterile technique!
2. Grow OP50 *E. coli* overnight shaking at 30°C (150-200 rpm). ~30ml in 50ml falcon tube.
3. Grow *L. musarum* overnight shaking at 25°C (50-150rpm). 5-6ml LB in a 15ml falcon tube, arrange the tubes in the shaking incubator on their sides, instead of upright, to increase aeration.
4. To check for contamination in OP50, streak overnight cultures on XLD plates and grow overnight at 30°C. To check for contamination in *L. musarum* cultures, streak overnight cultures on LB plates and grow for 48 hours at 25°C. *L. musarum* colonies are well defined, small, yellowish, and 'sticky'.

Freezing bacteria

1. 500µl of 50% glycerol into each cryotube
2. Add 500µl of overnight bacterial culture into each cryotube then invert tubes to mix
3. Store samples in -80°C freezer.

To 'seed' NGM with E. coli OP50

1. Pipette 100-200µl of OP50 culture onto 9cm NGM plate.
2. Use a L-shaped plate spreader to spread the bacteria.
3. Grow lawns at 30°C overnight.

Making experimental parasite 'exposure' plates

For experimental evolution we used 90mm plates, and for assays we used 60mm and 35mm plates.

1. Grow OP50 and *L. musarum* cultures in liquid culture overnight as described above. Worms don't eat *L. musarum*, so we create a mix *L. musarum* and OP50 when exposing worms.
2. Measure the $OD_{630}$ of all OP50 and *L. musarum* cultures, and dilute:
    a. *L. musarum* = $OD_{630}$ 0.2
    b. OP50 = $OD_{630}$ 0.4

To calculate volumes needed to dilute use the formula C1V1 = C2V2. For 6cm exposure plates, you will want about 100µl of bacterial culture per plate, so work out what final volume of bacteria you will need to seed all exposure plates.

3. Create a *L. musarum*:OP50 mixture with your diluted cultures, with 20% *L. musarum* and 80% OP50
4. Vortex *L. musarum*:OP50 cocktail to mix (it is ok to vortex *L. musarum* once it is very dilute)
5. Pipette 200µl (90mm plates), 100µl (60mm plates), or 50µl (35 plates) of the *L. musarum*:OP50 mixture onto each NGM plate and spread with a spreader covering most of the plate
6. Incubate overnight at 25°C

Washing worms off plates

1. Add 5ml M9-Tx to a worm plate.
2. Swill the liquid around and gently use L-shaped spreader to dislodge the worms.
3. If you are not worried about disturbing the bacterial lawn or if you are trying to also get eggs use the pipette to "pressure wash" the worms off (they can stick to the plate).
4. Tilt the plate to allow worms to sink to one edge, then pipette wormy M9 into a labelled falcon tube (careful not to suck up the agar)

Bleaching worms

Only bleach plates where you can see eggs and gravid hermaphrodites (ones with eggs in them) to ensure success!

Wash worms off your plates using M9 (see "washing worms off plates"). Use a sterile L-shaped spreader to scrape the surface of plates to dislodge eggs.

Pipette wormy M9 into 14ml falcon tubes so that there is 4ml in each tube.

I sometimes combine worms from multiple plates into single falcons (3 plates max).

If you have washed multiple plates into each falcon tube, centrifuge the tubes at 3500 for 2 minutes, and pipette off supernatant, leaving 4ml in each tube.

In a separate falcon tube, mix 50:50 ratio of bleach (10-12% sodium hypochlorite) and 5M NaOH.

You want enough 50:50 bleach NaOH mixture to be able to add at least 1ml to each of your worm falcon tubes.

Make sure the sodium hypochlorite is fresh (it degrades in light over weeks) but also not too strong. Dilute bleach 50:50 with water or M9 if it is too strong.

Add 1ml of the bleach NaOH solution to each 4ml worm sample.

Shake/vortex regularly for 7-8 minutes.

At first, all the worms go really straight, like sticks. Then they usually tend to bend in the middle (in hermaphrodites, this is where the vulva is and so is quite easily penetrated by the solution). The liquid in the tube should also become less cloudy. If it does not the bleach may be too weak.

After 7-8 minutes vortex the tubes for up to 1 minute.

Place the falcon tubes in the centrifuge and spin for 2 minutes at 3500rpm.

Remove the supernatant (leaving the pellet). Pipette in 5ml M9. Centrifuge again

Remove the supernatant (leaving the pellet). Pipette in 5ml M9. Centrifuge again

After this second wash, remove the supernatant (leaving the pellet). Pipette 3ml M9 into each tube and vortex to re-suspend the pellet.

To achieve L1 arrest shake put the tubes overnight in the 20°C.

Some research suggests long periods in M9 is not great for worms, however.

<u>Plating</u> <u>L1's</u> <u>on</u> <u>OP50</u>

Take tubes from 20°C.

If your wormy M9 has lots of dead animals (after bleaching):

allow the sample to stand in a falcon rack for a minute or so to allow most of the large dead worms to settled to the bottom. Don't wait too long or too many L1's will settle as well.

After a minute or so, pipette most of the supernatant into a new falcon, leaving the majority of the dead worms in the old falcon.

Count the number of worms per µl of your sample:

Briefly vortex the sample so that all the worms are homogenous through the sample.

Pipette six 2µl droplets of the sample onto an empty petri dish. Count the number of worms present under the microscope.

Calculate the average of the number of worms per µl.

Calculate the number of µls you need to put onto each plate (this depends on the amount of worms you want on each plate. 1000-2000 L1's per 9cm plate).

E.g. I want to plate 1000 L1 worms and I counted 10 worms per µl: 1000/10 = 100µl per plate.

It is good to avoid plating high volumes of liquid. If there are very few L1's per µl, they can be concentrated by centrifuging at 1600rpm for 1 minute. Remove some supernatant, vortex and then re-count.

At this point you can work out whether you have enough worms for your experiment by looking at the total volume of wormy M9 you have.

Pipette calculated volume onto NGM plates seeded with OP50 (see "To seed NGM"). Between plates regularly vortex/shake sample to keep the worms distributed evenly in the sample.

Allow the plates to dry in the biological safety cabinet and put them in the incubator of choice (usually 20°C if growing worms for an experiment).

Exposing worms to parasite

Wash worms off your OP50 plates using M9 (see "washing worms off plates").

If you expect that there will be a low number of worms per µl, concentrate sample by centrifuging at 1600rpm for 1 minute, and remove a some of the supernatant.

Count the number of worms per µl of your sample:

Briefly vortex the sample so that all the worms are homogenous through the sample.

Pipette six 2µl droplets of the sample onto an empty petri dish. Count the number of worms present under the microscope.

Calculate the average of the number of worms per µl.

Calculate the number of µls you need to put onto each plate (this depends on the amount of worms you want on each plate). Re-concentrate worms if needed to plate the smallest volume that is reasonable.

E.g. I want to expose 100 worms per plate and I counted four worms per µl: 100/4 = 25µl per plate.

At this point you can work out whether you have enough worms for your experiment by looking at the total volume of wormy M9 you have.

Pipette this amount of the sample onto each plate that you have previously prepared (see "making exposure plates"). Do this whilst continually shaking the sample to keep the worms distributed evenly in the sample.

Allow the plates to dry in the biological safety cabinet and put them in the incubator of choice (this depends on your experiment).

If there are still dead worms in your sample from the bleaching process use a marker pen to circle around the area where you spotted the worms. When you come to count mortality you can avoid counting any worms within this area to avoid counting worms that died prior to the assay. If the volume plated is as small as possible, the area that needs to be avoided should also be small.

Counting mortality

It's best to count mortality without knowing the treatments you are counting- code the plates so you don't unknowingly bias your data.

Draw lines on the bottom of the plate or the lid of a spare plate, splitting the plate into sections of 4/5.

Go across the plate and count the total number of worms visible (dead and alive) using a hand clicker. Note this down. Use the segments on the plate as guides- count one section before moving on to the next). Doing it at reasonable speed is best as worms move!

Go back through each section and now count the number of dead worms. Poke them with a platinum wire to see if they move if you are unsure if they are alive or not.

Divide the number of dead by the total number of worms.

Washing worms

Use a platinum wire to pick up the worms from exposure plates and place them into a 1.5ml Eppendorf tube filled with 1200µl M9 with Triton-x. Check the pick carefully to make sure all worms have gone into the tube.

Centrifuge tube(s) at 2000rpm for 1 minute.

Remove 1000µl of the supernatant, being careful not to suck up any worms.

Pipette 1000µl M9-X into tubes.

Centrifuge at 2000rpm for 1 minute

Remove 1000µl of the supernatant, being careful not to suck up any worms.

Pipette 1000µl M9-X into tubes.

Centrifuge at 2000rpm for 1 minute

Remove 1000µl of the supernatant, being careful not to suck up any worms.

Pipette 1000µl M9 (without Triton-x) into tubes.

Centrifuge at 2000rpm for 1 minute

Remove 1000µl of the supernatant, being careful not to suck up any worms.

The worms will now be reasonably clean

Colony forming units (CFU) of bacteria in worms

Prepare one Eppendorf tube (2ml if using Qiagen Tissue Lyser) per replicate with exactly 1200µl M9-x and pour in around 10 sterile beads. Be as sterile as possible.

Under the microscope pick 10 worms per replicate (i.e. plate) into the Eppis.

Do this with a flame on to keep the Eppis sterile while they are open and to flame the picker. Check the pick under the microscope to make sure all the worms have been transferred to the M9. Flame the picker between each replicate to sterilise.

Once all replicates are picked, wash them according to "Eppendorf" method (see above).

After the final wash step you want exactly 200µl of M9 along with your worms left in the tubes.

Crush the worms with a tissue lyser.

Bead beater settings: Speed depends on which one you use, but do max speed for 3-8 minutes. After crushing the worms, hold the tubes up to a light and gently flick the tubes – you will be able to see worms floating about if the crush hasn't worked.

Create a dilution series of your crushed sample in a 96-well plate using M9 that does NOT contain triton-x.

Use a multichannel pipette to put 180µl of M9 into each well.

In the first row of wells, pipette 20µl of each neat sample (this first well will be a 10-1 dilution).

Use the multichannel pipette to then make a serial dilution down to 10-4 by pipetting 20µl of each dilution into the subsequent one. Mix each dilution by aspirating the pipette up and down 3 times as you dispense the sample.

Change tips between each dilution.

Plate between three and six 10-20µl spots of multiple dilutions (for *L. musarum* 10-3 & 10-4 are often best) on 90mm LB plates. Make sure plates are labelled well so you know which the sample ID and dilution of each set of spots. You should be able to fit between two and four samples/dilutioons per 90mm LB plate.

Allow to dry in the sterile cabinet, then put plates at 25°C for 48 hours or until you can see distinct colonies.

Once colonies are visible, they can be counted straight away, or plates can be put at 4°C and counted later.

Use a clicker or colony counter to count the number of colonies. Using a marker pen you can mark each colony as you count to keep track. Don't forget to calculate CFU per worm from your colony counts!

Equation:

CFU per worm=((colony count*dilution factor)*((initial sample volume)/(plating volume)))/(worm number)

E.g. CFU per worm=((colony count*10^(-4) )*(200ul/(20ul*6 spots)))/(10 worms)
